# Microtubule-mediated GLUT4 trafficking is disrupted in insulin resistant skeletal muscle

**DOI:** 10.1101/2022.09.19.508621

**Authors:** Jonas R. Knudsen, Kaspar W. Persson, Carlos Henriquez-Olguin, Zhencheng Li, Nicolas Di Leo, Steffen H. Raun, Janne R. Hingst, Raphaël Trouillon, Martin Wohlwend, Jørgen F. P. Wojtaszewski, Martin A. M. Gijs, Thomas E. Jensen

## Abstract

Microtubules serve as tracks for long-range intracellular trafficking of glucose transporter 4 (GLUT4), but the role of this process in skeletal muscle and insulin resistance is unclear. Here, we used fixed and live-cell imaging to study microtubule-based GLUT4 trafficking in human and mouse muscle fibers and L6 rat muscle cells. We found GLUT4 localized along and on the microtubules in mouse and human muscle fibers. Pharmacological microtubule disruption using Nocodazole (Noco) prevented long-range GLUT4 trafficking and depleted GLUT4-enriched structures at microtubule nucleation sites in a fully reversible manner. Using a perfused muscle-on-a-chip system to enable real-time glucose uptake measurements in isolated mouse skeletal muscle fibers, we observed that Noco maximally disrupted the microtubule network after 5 min without affecting insulin-stimulated glucose uptake. In contrast, a 2h Noco treatment markedly decreased insulin responsiveness of glucose uptake. Insulin resistance in mouse muscle fibers induced either *in vitro* by C2 ceramides or *in vivo* by diet-induced obesity, impaired microtubule-based GLUT4 trafficking. In L6 muscle cells, pharmacological activation of the microtubule motor protein kinesin-1 increased basal and insulin-stimulated GLUT4 translocation, whereas shRNA-mediated knockdown of the kinesin-1 protein encoding gene *Kif5B* reduced insulin-stimulated GLUT4 translocation. Thus, in adult skeletal muscle fibers, the microtubule network is essential for intramyocellular GLUT4 movement, likely functioning to maintain an insulin-responsive cell-surface recruitable GLUT4 pool via kinesin-1 mediated trafficking.

## Introduction

Skeletal muscle is quantitatively the largest site of glucose disposal, a process facilitated by insulin and contraction-responsive translocation and insertion of glucose transporter 4 (GLUT4) into the surface membrane of muscle fibers (1, 2). Insulin resistant human and rodent muscle exhibit impaired insulin-stimulated GLUT4 translocation (3–6) and muscle-specific deletion of GLUT4 is sufficient to cause systemic insulin resistance and glucose intolerance (7). However, the details of GLUT4 regulation – particularly in adult skeletal muscle – and the causes of skeletal muscle insulin resistance remain unclear. In L6 myoblasts and 3T3-L1 adipocytes, insulin resistance not only decreases insulin-stimulated GLUT4 recruitment to the surface membrane, but also affects the distribution of GLUT4 between intracellular compartments (8, 9). This suggests that disturbed intracellular sorting of GLUT4 contributes to peripheral insulin resistance.

Motor protein-mediated trafficking on the microtubule cytoskeleton is well-established to allow long-range transport of a diverse assortment of molecules and to position intracellular organelles and membrane-structures in various cell types (10). For GLUT4, long-range microtubule-dependent GLUT4 movement beneath the plasma membrane has been observed in adipocyte cell culture (11) and similar long-range movement was also seen in adult rodent skeletal muscle (12). A requirement for microtubule-based protein trafficking is supported by several observations in cultured cells. Microtubule disruption dispersed perinuclear GLUT4 in 3T3-L1 adipocytes (13, 14) as well as L6 myoblasts (8) and impaired GLUT4 membrane insertion in some (8,14–16) but not all studies (17, 18). Whether microtubules are required for intracellular GLUT4 positioning and trafficking in adult skeletal muscle is unknown, nor has the influence of insulin resistance on microtubules and/or microtubule-based GLUT4 trafficking been investigated in skeletal muscle fibers.

Therefore, we here characterized various aspects of microtubule-based GLUT4 trafficking in adult human and mouse skeletal muscle, supplemented with mechanistic studies in cultured L6 muscle cells. Our findings suggest that an intact microtubule network is required for KIF5B-mediated intracellular GLUT4 movement and maintaining insulin-responsive glucose uptake, whereas skeletal muscle insulin resistance leads to impaired microtubule-based GLUT4 trafficking.

## Results

### GLUT4 was enriched at microtubule nucleation sites and travelled on microtubule filaments in adult mouse and human muscle

To study the involvement of microtubules in GLUT4 trafficking, we first used structured illumination microscopy to image the subsarcolemmal (up to 4 µm into the muscle fiber) microtubule network and GLUT4 in mouse and human skeletal muscle at super resolution. In both mouse flexor digitorum brevis (FDB) (Figure 1A) and human vastus lateralis (Figure 1B) muscle, we observed GLUT4 localized on microtubule filaments and was enriched at microtubule filament intersections, previously identified as microtubule nucleation sites (19). Next, to study GLUT4 live, we overexpressed GLUT4-7myc-GFP (GLUT4-GFP) (20) alone or together with mCherry-Tubulin in mouse FDB muscle fibers (Figure 1C). GLUT4-GFP was localized in the same pattern as endogenous GLUT4 and found along the microtubule network, including on the more stable subpopulation (21) of detyrosinated microtubules (Figure 1 – figure supplement 1A) which has been implicated in trafficking of lipid droplets, mitochondria and autophagosomes in other cells types (22, 23) as well as on mCherry-Tubulin labelled microtubules (Figure 1 – figure supplement 1B). Live-imaging revealed long-range lateral directional movement of GLUT4 along filamentous tracks (Figure 1 – figure supplement 1C), corresponding to mCherry-Tubulin containing microtubule filaments (Figure 1D and Figure 1 – figure supplement 2). The GLUT4 structures occasionally exhibited long tubular morphology (>2 µm) but were mostly minor tubular structures or spheres (size varying from ∼0.4 μm^2^ down to the unresolvable) observed to undergo budding and fusion events on the microtubule tracks (Figure 1 – figure supplement 1D and E). Live-imaging, including fluorescence recovery after photobleaching (FRAP) experiments, revealed particularly dynamic and bidirectional movement at the microtubule nucleation sites (Figure 1 – figure supplement 1F-H and Figure 1 – figure supplement 3). Collectively, a portion of GLUT4 localized to microtubule nucleation sites and on microtubule filaments in adult mouse and human skeletal muscle. Furthermore, GLUT4 underwent continuous movement, budding and fusion along the microtubule tracks in live mouse skeletal muscle.

**Figure 1:**
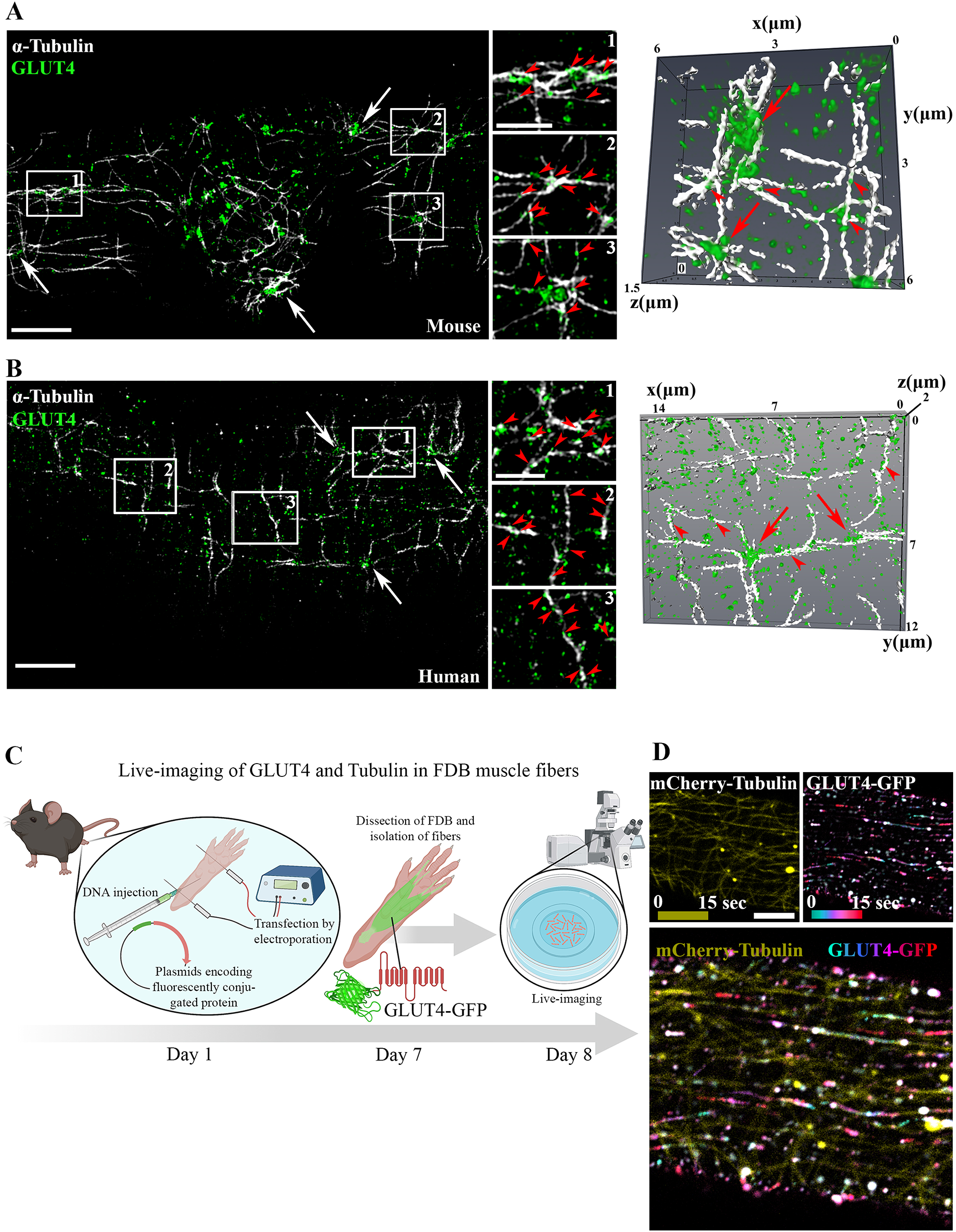
GLUT4 was enriched at microtubule nucleation sites and travelled on microtubule filaments in mouse and human muscle. A-B) Structured illumination microscopy (SIM) imaging of endogenous α-tubulin and GLUT4 (left panel) and 3D reconstruction of GLUT4 (green) and α-Tubulin (white) (right panel) in mouse flexor digitorum brevis (FDB) muscle (A) and human vastus lateralis muscle (B). Arrows indicate GLUT4 at microtubule nucleation sites, arrowheads mark GLUT4 vesicles along the microtubule filaments. C) Overview of workflow for live-imaging of fluorescently conjugated proteins in adult mouse FDB muscle fibers. D) Live-imaging of FDB expressing GLUT4-GFP and mCherry-Tubulin. Yellow projection of mCherry-Tubulin outlines the microtubule filaments (top panel left). Movement of GLUT4-GFP was visualized by color coded projection (first image cyan, last image red, top panel right). The merged projection (bottom), demonstrated movement of GLUT4-GFP along the mCherry-Tubulin containing microtubule filaments indicated by color-coded projections on top of the microtubule filaments. The movement of GLUT4-GFP is shown in Movie 1. A-B). Images are representative of >5 fibers from ≥3 different mice in A+D and 3 different fibers from 3 different subjects in B. Scale bar = 5 µm (A, B, D) and 2 µm (inserts in B, D).

### GLUT4 trafficking and localization required intact microtubules

Next, we tested if microtubule-based GLUT4 trafficking was insulin-responsive and dependent on an intact microtubule network. Insulin (30 nM) stimulation increased insulin signaling at the level of Akt Thr308 as expected (Figure 2 – figure supplement 1A), and the microtubule depolymerizing compound Nocodazole (Noco) (13 µM) significantly reduced both the total and the Noco-resistant (24) detyrosinated pool of polymerized microtubules by ∼90% and ∼50%, respectively (Figure 2 – figure supplement 1B and C). We observed a ∼16% but insignificant increase in GLUT4 movement on microtubules upon insulin stimulation and this movement was completely prevented by microtubule depolymerization (Figure 2 A and B, Figure 2 – figure supplement 1D, Figure 2 – figure supplement 2). This modest effect of insulin was observed across all insulin-stimulation experiments in the current study and averaged ∼20% (Figure 2C). Having established that GLUT4 trafficking was dependent on the microtubule network, we next tested if microtubule disruption affected the overall GLUT4 localization and distribution between different compartments. For quantification, we divided the GLUT4 structures into size categories corresponding to a) large GLUT4-positive structures at the microtubule nucleation sites (> 4 µm^2^), b) intermediate sized endomembrane structures (0.4-4 µm^2^) (25, 26)) and c) the smallest resolvable endomembrane structures (<0.4 µm^2^) including presumably insulin-responsive GLUT4 storage vesicles (GSVs) (Figure 2 – figure supplement 1E). Microtubule disruption by Noco (13 µM) drained the GLUT4 structures at the microtubule nucleation sites and reduced the amount of the smallest structures, while causing an increase in the intermediate sized structures (Figure 2D and E). These changes were reversed within 9 hours after removal of Noco (Figure 2D and E). Within the smallest structures, there was a shift toward fewer GLUT4 containing membrane structures, but each structure with a larger area (Figure 2 – figure supplement 1F). The total number and area of GLUT4 structures did not differ between conditions (Figure 2 – figure supplement 1G). In a previous study in L6 myoblasts, microtubule disruption prevented pre-internalized GLUT4 from reaching a Syntaxin6-positive perinuclear sub-compartment involved in GSV biogenesis and from undergoing insulin-responsive exocytosis (8). We therefore tested in adult muscle, if microtubule disruption similarly prevented accumulation in a Syntaxin6 sub-compartment. However, a limited and Noco-insensitive (in mice) co-localization was observed between Syntaxin6 and endogenous GLUT4 in human and mouse skeletal muscle, and fluorescent GLUT4-EOS (27) in mouse skeletal muscle (Figure 2F and G, Figure 2 – figure supplement 1H). Altogether, our data demonstrate that insulin has a modest stimulatory effect on the number of GLUT4 moving on microtubule and that GLUT4 trafficking and distribution is disrupted by pharmacological microtubule network depolymerization in a fully reversible manner.

**Figure 2:**
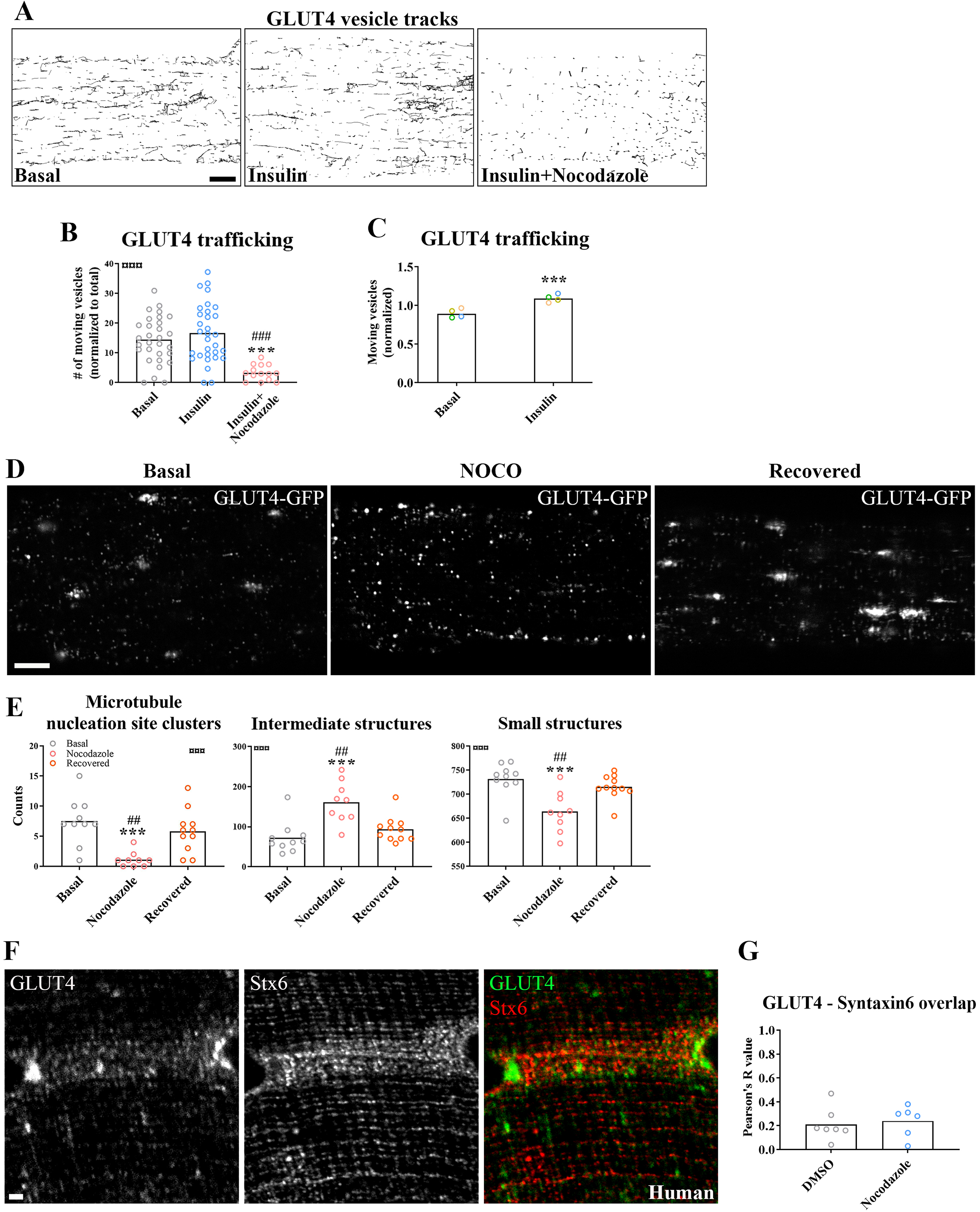
GLUT4 trafficking and localization was dependent on an intact microtubule network. A) Representative time-lapse traces of GLUT4-GFP vesicle tracking in muscle fibers ± insulin (INS, 30 nM) for 15-30 minutes with or without microtubule network disruption by Nocodazole (Noco, 13 µM) for 4 hours prior to insulin. The dynamics of GLUT4-GFP in the different conditions are also exemplified in movie 3. B) Quantified microtubule-based GLUT4 trafficking. C) Quantification of microtubule-based GLUT4 trafficking in basal and insulin-stimulated fibers pooled from 4 independent experiments. D) Representative images of muscle fibers ± pre-treatment with Noco (13 µM, for 15 hours, followed by recovery in Noco-free medium for 9 hours. E) Quantification of GLUT4 distribution between the microtubule nucleation sites (structures sized > 4 µm2), intermediate sized structures (0.4-4 µm2) and the smallest resolvable structures (<0.4 µm2) in fibers treated as in D. Compartment identification is described in figure S2 E. F) GLUT4 and Syntaxin6 (Stx6) in muscle fiber from human vastus lateralis muscle. G) GLUT4-Stx6 overlap in mouse flexor digitorum brevis muscle fibers in DMSO medium with and without Noco (13 µM) treatment. For A-B, n ≥ 14 muscle fibers from 5 different mice. For D-E, n = 9-11 muscle fibers from 3 different mice. For F, n= 3 subjects. Data are presented as mean with individual data points. *** p<0.001 different from basal, ### p<0.001 different from INS, ## p<0.01 different from Noco recovery. ¤¤¤ p<0.001 ANOVA effect Scale bar = 5 µm (A-D) and 2 µm (F).

### Prolonged, but not short-term, microtubule disruption blocked insulin-induced muscle glucose uptake

To investigate the requirement of microtubule-based GLUT4 trafficking and localization for insulin-induced muscle glucose uptake, we assessed muscle glucose uptake ± insulin and ± microtubule disruption in isolated incubated whole mouse soleus and extensor digitorum longus (EDL) muscles. When mouse soleus and EDL muscles were incubated *ex vivo* ± insulin and ± Noco (13 µM) for 15 min and up to 2h, an interaction between insulin and Noco was observed and the insulin-induced glucose uptake was gradually impaired over time and completely disrupted after 2h in both muscles (Figure 3A). The increase by insulin stimulation was significantly impaired after 40 min in soleus and 2h in EDL (Figure 3B). Insulin-stimulated signaling via p-Akt and p-TBC1D4 was unaffected by Noco treatment (Figure 3 – figure supplement 1A-C).

**Figure 3:**
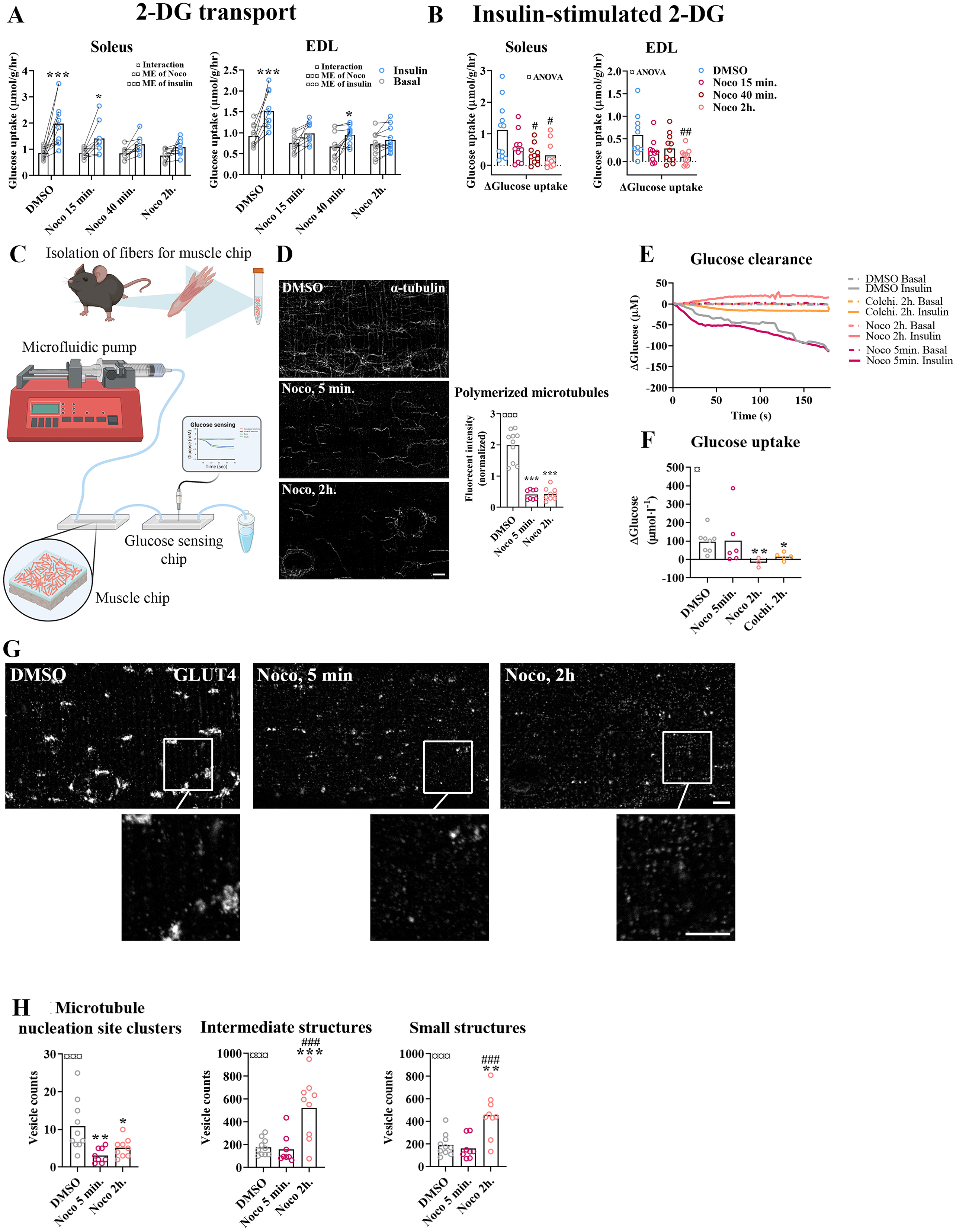
Time-dependent effect of microtubule disruption on insulin-induced muscle glucose uptake. *A)* 2-Deoxyglucose (2-DG) transport in basal and insulin-stimulated mouse soleus and extensor digitorum longus (EDL) muscles pretreated with Nocodazole (Noco, 13 µM) for the indicated time. *B)* Insulin-stimulated 2-DG transport (insulin minus basal) from muscles shown in A. C) Experimental setup for muscle on a chip system with glucose sensor. D) Microtubules imaged with α-tubulin in GLUT4-GFP expressing mouse flexor digitorum brevis (FDB) fibers treated ± Noco (13 µM) for 5 min or 2h. E) 180 sec. measurements of glucose concentration in perifusate from basal and insulin-treated FDB muscle fibers in muscle chips pre-incubated with DMSO, Noco (13 µM, 5min or 2h) or colchicine (25 µM, 2h). F) Insulin-stimulated glucose uptake into FDB muscle fibers in muscle chips calculated from the last 20 seconds of the concentration curves in E. G) Representative GLUT4 images from isolated mouse FDB muscle fibers treated ± Noco (13 µM) for 5 min. or 2h. H) Quantification of GLUT4 in large, intermediate and small sized membrane structures in FDB fibers treated ± Noco (13 µM) for 5 min. or 2h. The membrane compartment division by size is shown in figure S2 E. For A-B n = 6-7 muscles from 6-7 mice. For D and G-H, n =8-10 muscle fibers from 3 different mice. For E-F, n ≥ 3 muscle chips from 3-4 mice. Data are presented as mean with individual data points. Paired observations from the same mouse are indicated by a connecting line. */**/*** p<0.05/0.01/0.001 different from basal/DMSO, #/## p<0.05/0.01 different from DMSO. ¤/¤¤/¤¤¤ p<0.05/0.01/0.001 ANOVA effect. Scale bar = 5 µm.

To understand the temporal resolution of microtubule network disruption and its effect on insulin-induced glucose uptake, we investigated adult isolated single fibers at much higher time resolution, using an adopted chip system (28, 29) with a customized glucose sensing electrode for glucose uptake measurements (Figure 3C) and characterized its feasibility to measure glucose uptake in skeletal muscle fibers (Figure 3 – figure supplement 1D-F). This system measures continuous glucose concentration in perifusate post muscle fiber exposure, and hence glucose uptake in real-time. This method allowed us to estimate glucose uptake in isolated FDB fibers at ∼5 µM glucose concentration sensitivity (defined as a registered fluctuation of thrice the standard deviation of the baseline measurements) with a temporal resolution of <4 sec. (Figure 3 – figure supplement 1G and H). Noco (13 µM) caused complete microtubule disruption in FDB fibers within 5 minutes similar to a 2-hour treatment (Figure 3D). Interestingly, acute microtubule disruption (5 min Noco) affected neither basal (Figure 3 – figure supplement 1I) nor insulin-induced muscle glucose uptake, whereas 2 hours treatment by Noco or colchicine (25 µM), another microtubule disrupter, completely blocked insulin-induced muscle glucose uptake (Figure 3E and F). Notably, GLUT4 containing large membrane structures corresponding mainly to microtubule nucleation sites were already reduced after 5 min of Noco exposure, whereas accumulation of GLUT4 in intermediate and small sized membrane structures was only observed after 2h of Noco exposure (Figure 3G and H). Thus, an intact microtubule network is not required for the immediate insulin-induced GLUT4 translocation response in adult skeletal muscle fibers. However, prolonged disruption of the microtubule network causes a more pronounced missorting of GLUT4 and renders skeletal muscle unresponsive to insulin.

### KIF5B-containing kinesin-1 motor proteins regulate muscle GLUT4 trafficking

Next we investigated which motor protein(s) mediate microtubule-dependent GLUT4 trafficking in skeletal muscle. Since the kinesin-1 heavy chain protein encoding gene *Kif5b* has been implicated in GLUT4 trafficking in adipocytes (30, 31), we studied the consequence of Kinesin-1 microtubule motor protein family activation as well as knockdown of *Kif5b* using L6 skeletal muscle cells overexpressing exofacially tagged GLUT4 (32, 33) (Figure 4A). First, we assessed the GLUT4 surface content in L6 myoblasts following stimulation with the small-molecule Kinesin-1 activator Kinesore (34). Kinesore dose-dependently increased GLUT4 surface content up to 50 µM Kinesore (Figure 4 – figure supplement 1A). In addition to the increased GLUT4 surface content, Kinesore induced GLUT4 redistribution from the perinuclear compartment towards the cell periphery (Figure 4B). Similarly in isolated mouse FDB fibers, Kinesore decreased the GLUT4 content at the microtubule nucleation sites (Figure 4C). The effect of kinesore on GLUT4 surface content was additive to insulin (Figure 4D). No effect was observed on phosphorylation of AMPK, an insulin-independent stimulator of GLUT4 translocation, whereas a modest potentiation of insulin-stimulated Akt Thr308 but not Ser473 phosphorylation was observed (Figure 4 – figure supplement 1B). As expected, glucose transport into L6 myoblasts was also increased by Kinesore (Figure 4 – figure supplement 1C) as was GLUT4 membrane insertion in L6 myotubes (Figure 4E). Next, using a short hairpin (sh) construct, we lowered KIF5B expression by ∼70% in L6 myoblasts (Figure 4 – figure supplement 1D). This did not affect GLUT4 expression (Figure 4 – figure supplement 1E) but impaired insulin-stimulated GLUT4 translocation (Figure 4F). Collectively, these data strongly suggest the involvement of KIF5B-containing Kinesin-1 motor proteins in plus-end directed GLUT4 transport along microtubules in skeletal muscle.

**Figure 4:**
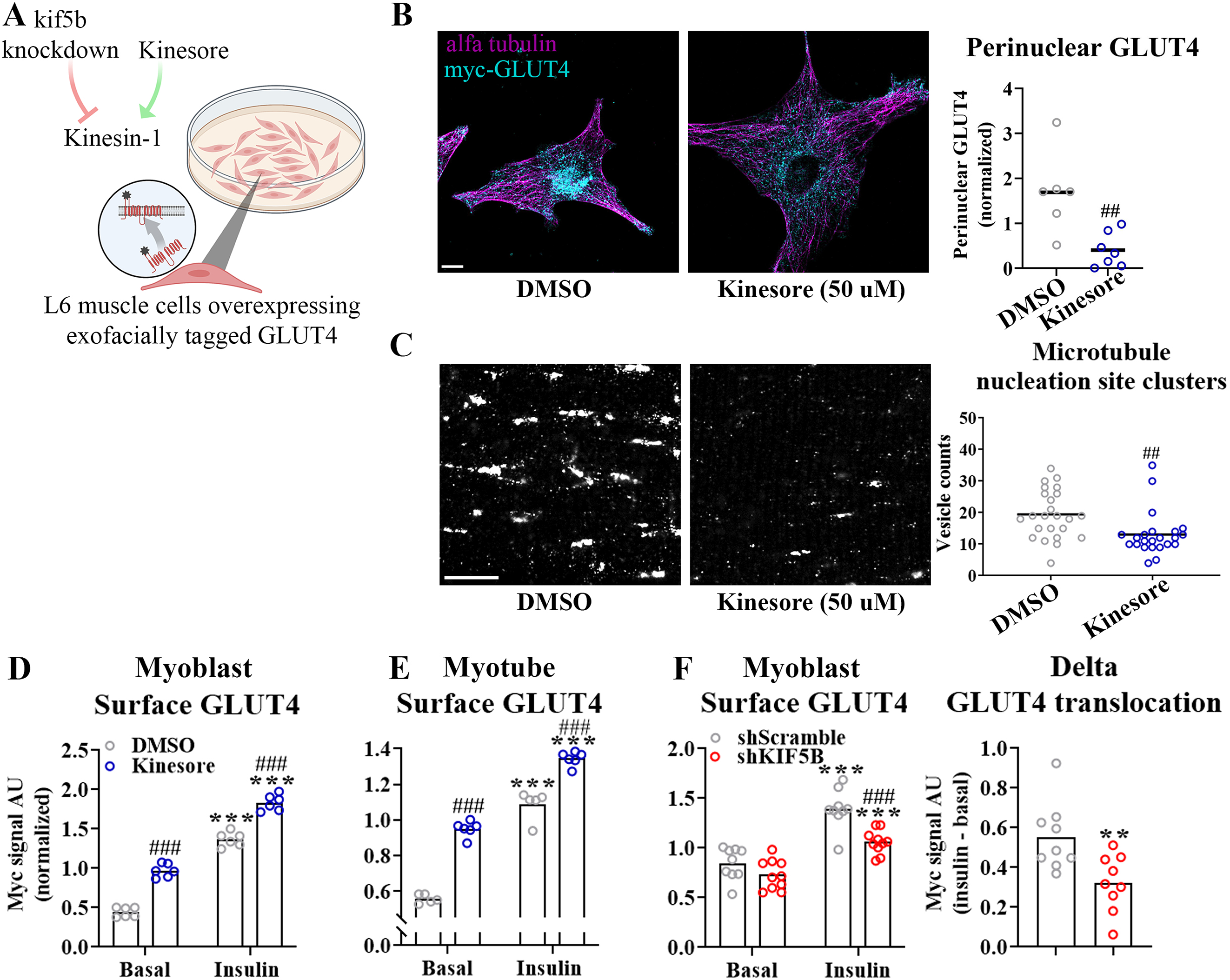
Kinesin-1 containing Kif5b regulated GLUT4 localization and translocation. *A)* Schematic overview of L6 muscle cell system to assess GLUT4 surface content. B) Imaging (left) of α-tubulin (magenta) and GLUT4 (cyan) and quantification (right) of perinuclear GLUT4 in L6 myoblasts ± kinesore treatment (2h, 50 µM). C) Imaging (left) of GLUT4 and quantification (right) of GLUT4 at the microtubule nucleation sites in mouse flexor digitorum brevis (FDB) muscle fibers treated ± kinesore (2h). D-E) Exofacial GLUT4 signal at the surface membrane in L6 myoblast (D) or 7 day differentiated myotubes (E) serum starved for 4h and treated ± kinesore (50 µM) for the last 2h before 15 min ± insulin (100 nM). ANOVA main effect of insulin (¤¤¤) and Kinesore (¤¤¤) and interaction (¤¤). F) Exofacial GLUT4 signal in serum starved (4h) basal and insulin-stimulated (100 nM, 15 min) L6 myoblasts (left) and insulin-response (insulin minus basal, right) in GLUT4 surface content. L6 myoblasts were transfected with short hairpin scramble RNA (shScramble) or shRNA targeting KIF5B 72h prior to the experiment. ANOVA main effect of insulin (¤¤¤) and shKIF5B (¤¤¤) and interaction (¤). B, n = 6-7 individual samples pooled from 2 independent experiments. C, n=23-24 muscle fibers from 2 different mice. D-F, each data point represents the average of 3 replicates and originate from at least 3 independent experiments. Data are presented as mean with individual data points. **/*** p<0.01/0.001 effect of insulin. ##/### p<0.01/0.001 different from DMSO/Scramble. ¤/¤¤/¤¤¤ p<0.05/0.01/0.001 ANOVA effect. Scale bar = 5 µm.

### Insulin resistance induced by C2 ceramide and high fat diet impaired microtubule-based GLUT4 trafficking

Having established an essential role for the microtubule network in GLUT4 trafficking and muscle glucose uptake, we proceeded to test if microtubule-based GLUT4 trafficking was impaired in insulin resistant states. We induced insulin resistance in adult mouse skeletal muscle both *in vitro using* short-chain C2 ceramide and *in vivo* using diet-induced obesity (Figure 5A). *In vitro*, treatment of isolated FDB muscle fibers with C2 ceramide (50 µM) impaired insulin-stimulated Akt Thr308 phosphorylation (Figure 5 – figure supplement 1A) and markedly reduced microtubule-based GLUT4 trafficking defined as the number of moving vesicles (Figure 5B) and the total microtubule-based travelling of GLUT4 structures (Figure 5 – figure supplement 1B). *In vivo*, mice fed a 60% high fat diet (HFD) for 10 weeks exhibited impaired insulin and glucose tolerance as well as reduced insulin-stimulated phosphorylation of Akt Thr308 and Akt substrate TBC1D4 Thr642 in isolated FDB fibers (Figure 5 – figure supplement 1C-E), confirming whole-body and skeletal muscle insulin resistance. Similar to C2 ceramide-treated fibers, HFD-exposed FDB muscle fibers exhibited impaired microtubule-based GLUT4 trafficking (Figure 5C, Figure 5 – figure supplement 1F). This prompted us to ask whether the microtubule polymerization was itself insulin-responsive and/or affected by insulin resistance. To test this, we transfected mouse FDB muscle fibers with the microtubule plus-end-binding protein EB3-GFP, which binds the tip of growing microtubules via its calponin homology domain and has previously been used for live-cell characterization of microtubule polymerization (35). As previously reported (19), EB3-GFP transfection allowed visualization of growing microtubules as a dynamic comet tail-like appearance, an effect completely prevented by the microtubule stabilizer taxol (10 µM) treatment (Figure 5 – figure supplement 1G, Figure 5 – figure supplement 2). In our data-sets, we analyzed the microtubule polymerization frequency (by counting EB3-GFP puncta (36)), the average polymerization distance, the total polymerization distance and the polymerization directionality following C2 ceramide exposure or HFD. Insulin tended (p=0.095) to increase the number of polymerizing microtubules by an average of 28% compared to basal fibers while C2 ceramide treatment reduced the amount of polymerizing microtubules significantly and taxol almost abolished microtubule polymerization (Figure 5D and E). C2 ceramide treatment also reduced the total polymerization distance while the average polymerization was unaffected (Figure 5 – figure supplement 1H).

**Figure 5:**
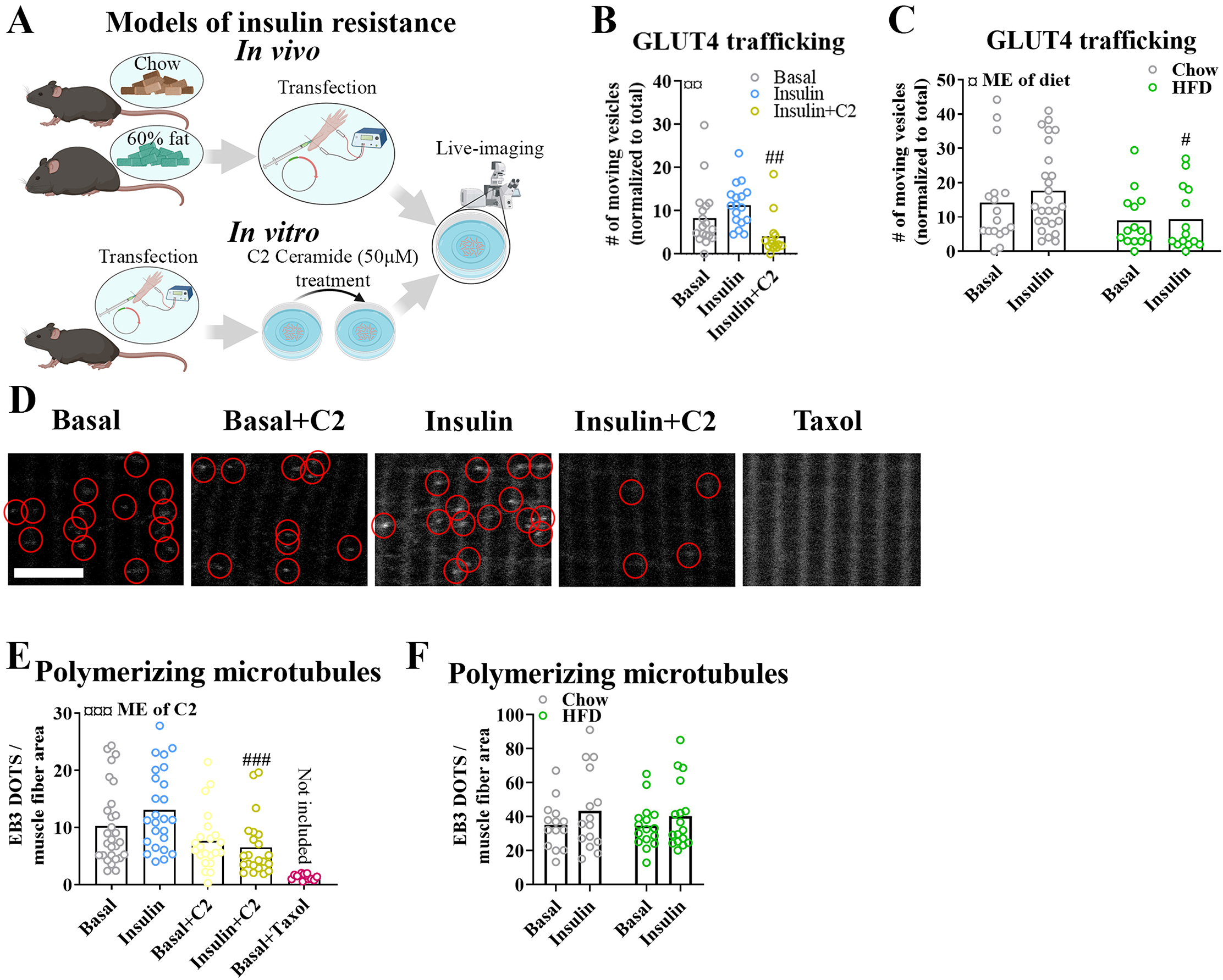
Insulin resistance impairs microtubule-based GLUT4 trafficking. *A)* Overview of in vitro and in vivo insulin resistance models used. B) Quantified microtubule-based GLUT4 trafficking in basal, insulin (INS, 30 nM) and insulin + C2 Ceramide (C2) (INS+C2, 30 nM+50 µM) treated flexor digitorum brevis (FDB) muscle fibers. C) Quantified microtubule-based GLUT4 trafficking in basal or INS (30 nM) treated FDB fibers from chow or high fat diet (HFD) fed mice. D) Representative images of polymerizing microtubules in EB3-GFP expressing FDB muscle fibers treated ± C2 (50 µM), paclitaxel (Taxol, 10 µM) for 2 hours prior to 15-30 minutes of INS (30 nM) stimulation. Red circles highlight microtubule tip-bound EB3-GFP. E) Quantification of polymerizing microtubules based on EB3-GFP in FDB fibers treated as in D. F) Quantification of polymerizing microtubules based on EB3-GFP in FDB fibers isolated from chow or 60% HFD fed mice and treated ± INS (30 nM) for 15-30 minutes. For B-F, n ≥ 13 muscle fibers from 3-4 mice. Taxol-treated muscle fibers were only used as a control and not included in the statistical analysis. NA = not statistically analysed. Data are presented as mean with individual data points. #/##/### p<0.05/0.01/0.001 different from INS (B) or different from corresponding group in chow fed mice (C) or control fibers (E). ¤/¤¤/¤¤¤ p<0.05/0.01/0.001 main effect (ME) of diet/C2.

In contrast to the effect of C2 ceramide on microtubule dynamics, HFD-induced insulin resistance was not associated with alterations in the number of polymerizing microtubules, average polymerization distance or total polymerization distance (Figure 5F, Figure 5 – figure supplement 1I). The polymerization directionality was also not affected by HFD (Figure 5 – figure supplement 1J). Altogether, different models of insulin resistance impaired microtubule-based GLUT4 trafficking in adult muscle fibers, suggesting a role in adult skeletal muscle insulin resistance. In contrast, defective microtubule polymerization was observed with C2 ceramide but not with the presumably more physiologically relevant HFD insulin resistance model.

## Discussion

In the present study, we provide translational evidence in adult human and mouse skeletal muscle, showing that the microtubule network is crucial for long-range directional GLUT4 trafficking via motor proteins, likely including KIF5B. Microtubule polymerization in isolated mouse muscle fibers could be abolished pharmacologically within minutes without affecting insulin-stimulated glucose uptake whereas longer pharmacological inhibition progressively caused GLUT4 mislocalization and lowered insulin responsiveness. These data are consistent with a model where microtubules are required for correct intramyocellular GLUT4 compartmentalization but not the ultimate insulin-stimulated GLUT4 translocation to the cell-surface from these compartments. Importantly, microtubule-based GLUT4 movement was impaired in two classical mouse insulin resistance models, short-chain ceramide treatment and diet-induced obesity. These data implicate dysregulation of microtubule-mediated GLUT4 trafficking and their localization in the etiology of adult skeletal muscle insulin resistance.

What may cause the reduced number of GLUT4 moving on microtubules in insulin resistant muscle? Based on cell-culture studies, GLUT4 is packaged into specialized GSVs, which upon insulin stimulation can undergo exocytosis (37). This exocytosis may include an intermediate step involving release and heterotypic fusion of GSVs with transferrin receptor positive endosomes observable using super-resolution quantum dot single GLUT4 particle tracking in isolated mouse soleus and EDL muscle fibers (38). Such model is consistent with previous reports of insulin-stimulated release of GSVs into an endosomal recycling pool in 3T3-L1 adipocytes (39, 40). In 3T3-L1 adipocytes, GSVs are synthesized via the trans-Golgi network (TGN) and the ER-Golgi intermediate compartment (41) by mechanisms involving TBC1D4 (42–44). We note that TBC1D4 phosphorylation on multiple sites is often decreased in insulin resistant skeletal muscle of humans (45, 46) and rodents (47, 48). Conversely, TBC1D4 phosphorylation is increased after insulin-sensitizing exercise/contraction (45,49–53) which might augment the insulin-responsive GLUT4 pool (54–56). The presently observed decreased GLUT4 movement on microtubules could reflect impaired availability of GLUT4 for trafficking, i.e. impaired TBC1D4-dependent GSV biosynthesis and/or heterotypic fusion of GSVs with transferrin receptor positive endosomes. Another possibility is that recruitment of GLUT4 onto microtubules, i.e. release of tethered GSVs and/or motor protein interaction, is impaired. In 3T3-L1 adipocytes, insulin stimulated the frequency but not movement speed of GLUT4 on microtubules, an effect blocked by a dominant-negative cargo-binding kinesin light chain 1 (KLC1) which impairs KIF5B function (30). Also, KLC1 was proposed to regulate GLUT4 translocation in 3T3-L1 adipocytes (30) and L6 myoblasts via insulin-regulated interaction with Double C2 domain B (57). These data are consistent with our observations of insulin-stimulated GLUT4 movement in adult mouse muscle and KIF5B dependence in L6 muscle cells. KIF5B was proposed to regulate GSV trafficking via insulin-stimulated binding to TUGUL, a cleavage product of the intracellular GSV tethering protein TUG (31, 58). Multiple studies by Jonathan Bogańs group linked TUG to skeletal muscle insulin-stimulated GLUT4 traffic and glucose uptake (59, 60). Insulin-stimulated TUG cleavage and expression of the putative protease Usp25m were reduced in rat adipose tissue after a 3-day high fat/high sugar diet-induced insulin resistance (31), suggesting possible dysregulation in insulin resistance. Apart from KIF5B and TUG, GLUT4 was proposed to utilize other motor proteins to move on microtubules in 3T3-L1 adipocytes, i.e. dynein via Rab5 (61), and the kinesin-2 family member KIF3A via AXIN and the ADP-ribosylase tankyrase 2 (62, 63). Whether any of these mechanisms regulate GLUT4 traffic in skeletal muscle should be tested further.

Our glucose uptake time-course data in perfused mouse FDB fibers suggest that microtubules are required to build and maintain the pool of insulin-responsive GLUT4 vesicles near the cell-surface observed in previous studies (12), and that disruption of this process may contribute to insulin resistant GLUT4 translocation in adult skeletal muscle. If true, then one would predict GLUT4 mislocalization to be observable in unstimulated insulin resistant skeletal muscle. Consistent with this prediction, Garvey and colleagues measured GLUT4 content in different sucrose density fractions in human vastus lateralis muscle biopsies and found a basal redistribution of GLUT4 to a denser fraction in type 2 diabetic subjects compared to control (6). Similar observations of altered intracellular GLUT4 distribution were made using a different fractionation protocol in subcutaneous adipose biopsies from type 2 diabetic subjects compared to control (64), implying that GLUT4 mislocalization is a shared hallmark of insulin resistant muscle and adipose cells. Application of higher resolution microscopy to the study of basal and insulin-stimulated GLUT4 localization in adult human and rodent insulin resistant skeletal muscle could help to resolve the relative distribution of GLUT4 between specific membrane compartments.

A previous study reported that the microtubule disrupting drug colchicine (25 µM for up to 8h) had no effect on insulin and contraction-stimulated glucose uptake, whereas Noco (3-83 µM for 30-60 min) potently inhibited glucose uptake in *ex vivo* incubated adult rat muscle and 33 µM Noco added for <5 min directly inhibited GLUT4 transporter activity into sarcolemma derived giant vesicles (65). Regarding colchicine, these divergent findings mirror previous studies in 3T3-L1 adipocytes where some support the absent effect of colchicine on insulin-stimulated cell surface GLUT4 translocation/glucose uptake despite strong microtubule disruption (17, 61) whereas others found colchicine to inhibit insulin-stimulated GLUT4 translocation (14, 16). The reason for this variation between studies is not readily apparent since the models and colchicine treatment protocols overlap. Regarding Noco, a direct inhibitory effect of 33 µM Noco added for >2 min on GLUT4 activity was suggested by previous 3T3-L1 adipocyte studies (17,18,66), However, Molero *et al.* also observed no inhibition of insulin-stimulated glucose uptake using 2 µM Noco for 1h despite complete disruption of microtubules (17), suggesting that Noco is useful to study the microtubule-glucose uptake connectivity at low concentrations. Since Noco was shown to inhibit GLUT4 activity within 2 min and we observed no effect of 13 µM Noco in isolated muscle fibers on basal glucose uptake or insulin-stimulated glucose uptake after 5 min at a concentration causing maximal inhibition of microtubule polymerization, we find it unlikely that Noco at the concentration used had major effects on GLUT4 activity.

When analyzed across our pooled control data-sets, insulin itself had a modest stimulatory effect on the number of GLUT4 moving on microtubules. This is consistent with live-muscle observations of GFP-tagged GLUT4 in rodent skeletal muscle where insulin increased the recovery of GLUT4 fluorescence after photo bleaching, suggesting that insulin increased the overall GLUT4 movement (67). Meanwhile, in primary and 3T3-L1 adipocyte cell culture total internal reflection fluorescence (TIRF) microscopy studies, insulin increased the number of GLUT4 halting and docking beneath the plasma membrane prior to insulin-stimulated insertion (11,68–70). TIRF imaging of mouse muscle fibers expressing HA-GLUT4-GFP suggested a similar GLUT4 halting and fusion-promoting effect of insulin in skeletal muscle (12). In regards to microtubule dynamics, although we did not detect any significant effect of insulin on the assessed parameters in adult muscle. Notably, insulin increased microtubule polymerization and/or density in 3T3-L1 adipocytes (71–73) and in L6 skeletal muscle cells (74) but was also observed to decelerate CLASP2 positive MT polymerization (71). Given that insulin may differentially increase or decrease the mobility of sub-populations of GLUT4 and microtubules, it seems likely that our relatively crude analyses of total subsarcolemmal GLUT4 movement could mask larger or even opposite effects on specific subpopulations.

In conclusion, we presently demonstrated that GLUT4 is present on microtubules in adult mouse and human skeletal muscle and that acute microtubule disruption causes intramyocellular GLUT4 redistribution and eventually decreases insulin-responsiveness of glucose transport. Decreased microtubule-dependent GLUT4 movement was observed in *in vitro* and *in vivo* mouse insulin resistance models, suggesting that disturbed microtubule-based GLUT4 trafficking is a hallmark of insulin resistance in adult skeletal muscle.

## Materials and methods

### Sample obtaining

*Human muscle samples* are tissue from *m. vastus lateralis* from young healthy men fasted for 6-7 hours. Further details on the subjects and tissue processing are described in previously published studies (55, 75).

*Mouse muscle samples* were from 10-16 weeks old C57BL/6 mice. All animal experiments were approved by the Danish Animal Experimental Inspectorate or by the local animal experimentation committee of the Canton de Vaud under license 2890 and complied with the European Union legislation as outlined by the European Directive 2010/63/EU. The current work adheres to the standards outlined in the ARRIVE reporting guidelines. Male C57BL/6NTac mice, 16 weeks old, were used for the experiments including high fat diet (HFD) fed mice. The mice were fed a 60% HFD or a control standard rodent chow diet *ad libitum*. For the rest of the experiments the mice were female C57BL/6JRj aged 10-16 weeks fed *ad libitum* with a standard rodent chow diet. All mice were exposed to a 12 h:12 h light-dark cycle.

### Gene transfer and fiber isolation

Flexor Digitorum Brevis (FDB) muscles were electroporated *in vivo* similar to (76) and isolated as previously described (77). The following plasmids were used: pB-GLUT4-7*myc*-GFP plasmid (a gift from Jonathan Bogan, Addgene plasmid #52872). p-mCherry-Tubulin-C1 (a gift from Kristien Zaal), HA-GLUT4-EOS (originally from the Zimmerberg laboratory (27) was a gift from Timothy McGraw) and p-EB3-GFP-N1 (originally from the Akhmanova laboratory (35) was a gift from Evelyn Ralston).

### Fiber culturing and drug treatments

Experiments with isolated fibers were performed the day after isolation. For prolonged (15 hours) nocodazole treatment, nocodazole (M1404, Merck) was added for a final concentration of 4 μg/ml at this step. When palmitic acid treatment is indicated, this was added for a final concentration of 0.5 mM at this step as well. Palmitic acid was dissolved to a 200 mM solution in 1:1 ethanol and α-mem, from which a 16x solution containing 100 mg/ml fatty acid free BSA was made. Non-treated fibers were treated with BSA without palmitic acid. When indicated C2 ceramide (50μM) (860502, Avanti) Paclitaxel (10 μM) (T7402, Merck), Kinesore (6664, Tocris) or Colchicine (25 μM) (C9754, Sigma) was added 2 hours prior to imaging/lysing whereas nocodazole (13 μM) was added 4 hours prior unless otherwise mentioned. For signaling analyses 30 nM insulin (Actrapid, Novo Nordisk A/S) was added 15 min prior to lysing, for microscopic analyses 30 nM insulin was added 15-30 min prior to imaging. For fixation fibers were incubated with 4% paraformaldehyde (Electron Microscopy Sciences) in PBS for 20 min.

### Cell culturing and experiments

L6 rat myoblasts expressing myc-tagged GLUT4 (33) were maintained in α-MEM (12561056, Gibco) supplemented with 10% fetal bovine serum and 1% pen/strep antibiotic in a humidified atmosphere containing 5% CO2 and 95% air at 37°C. Differentiation to myotubes were achieved by lowering the serum concentration to 2% for 7 days. For specific knock down of KIF5B shRNA constructs containing a 19-nucleotide (GGACAGATGAAGTATAAAT) sequence derived from mouse KIF5B mRNA (78) (A gift from Dr. Kwok-On Lai, City University of Hong Kong) were used using JetPRIME (Polyplus) according to manufacturer’s protocol. As control shRNA with the sequence CCTAAGGTTAAGTCGCCCTCGCTCGAGCGAGGGCGACTTAACCTTAGG, a gift from David Sabatini (Addgene plasmid # 1864) were used. Three days after initial transfection the experiments were conducted as described in the figure legends. For GLUT4 translocation assessment, cell surface GLUT4myc was detected using a colorimetric assay (79). Drug treatments were performed as described in the figure legends.

### Glucose uptake measurements

#### 2-DG transport

into Soleus and EDL muscles were assessed as described before (80).

#### 2-DG transport

into L6 cells was measured by washing cells in PBS containing HEPES and incubating them in PBS+HEPES containing 2-[3H] deoxyglucose for 5 min before cell harvest in lysis buffer. Tracer accumulation was then measured by liquid scintillation counting.

#### Electrochemical glucose sensing

Inspired by (81) we fabricated a microfluidic chip system using standard soft lithographic techniques. The chips were divided into a tissue chamber and a glucose sensing chamber connected by tubing. Both chambers were molded based on SU-8 master wafers. The tissue chamber consisted of 2 identical units each containing 3 layers of poly(dimethylsiloxane) (PDMS) (Figure 6A). The glucose sensing chamber was a single PDMS layer bonded to a glass slide by air plasma (Figure 6B). Fluid connection was achieved by punching holes using biopsy punchers and inserting tubes in the portholes (Figure 6C). A customized electrode was fabricated by threading a working electrode (platinum wire, Ø 51 µm, Teflon coated) and a reference electrode (silver wire, Ø 75 µm, Teflon coated) in the lumen of a 18 G syringe needle and embedding them in fluid epoxy. A counter electrode was attached to the metal of the needle by silver paste (Figure 6D). On the day of experiment the electrode was carefully polished using fine sand paper and alumina slury (0.05 µm particles) and a layer of chloride was deposited on the electrode by immersing it in 3 M KCl and exposing it to 6 current steps consisting of −20 µA for 1s followed by 20 µA for 9s. The working electrode was cleaned electrochemically in 0.1 M H_2_SO_4_ by running 10 cyclic voltammogram (CV) cycles. Next, to form an exclusion membrane on the censor, a layer of poly-(m-phenylenediamine) (mPD) was electropolymerized on to the working electrode by applying 20s of 0.0 V, 5 min at 0.7 volt and at 0.0 V. Finally the sensor was modified by addition of glucose oxidase by embedding a PBS solution consisting of glucose oxidase (60 mg*ml^-1^), bovine serum albumin (30 mg*ml^-1^) and poly(ethylene glycol) diglycidyl ether (60 mg*ml^-1^) and 2% glycerol on top of the electrode via 2h incubation at 50 °C.

**Figure 6.**
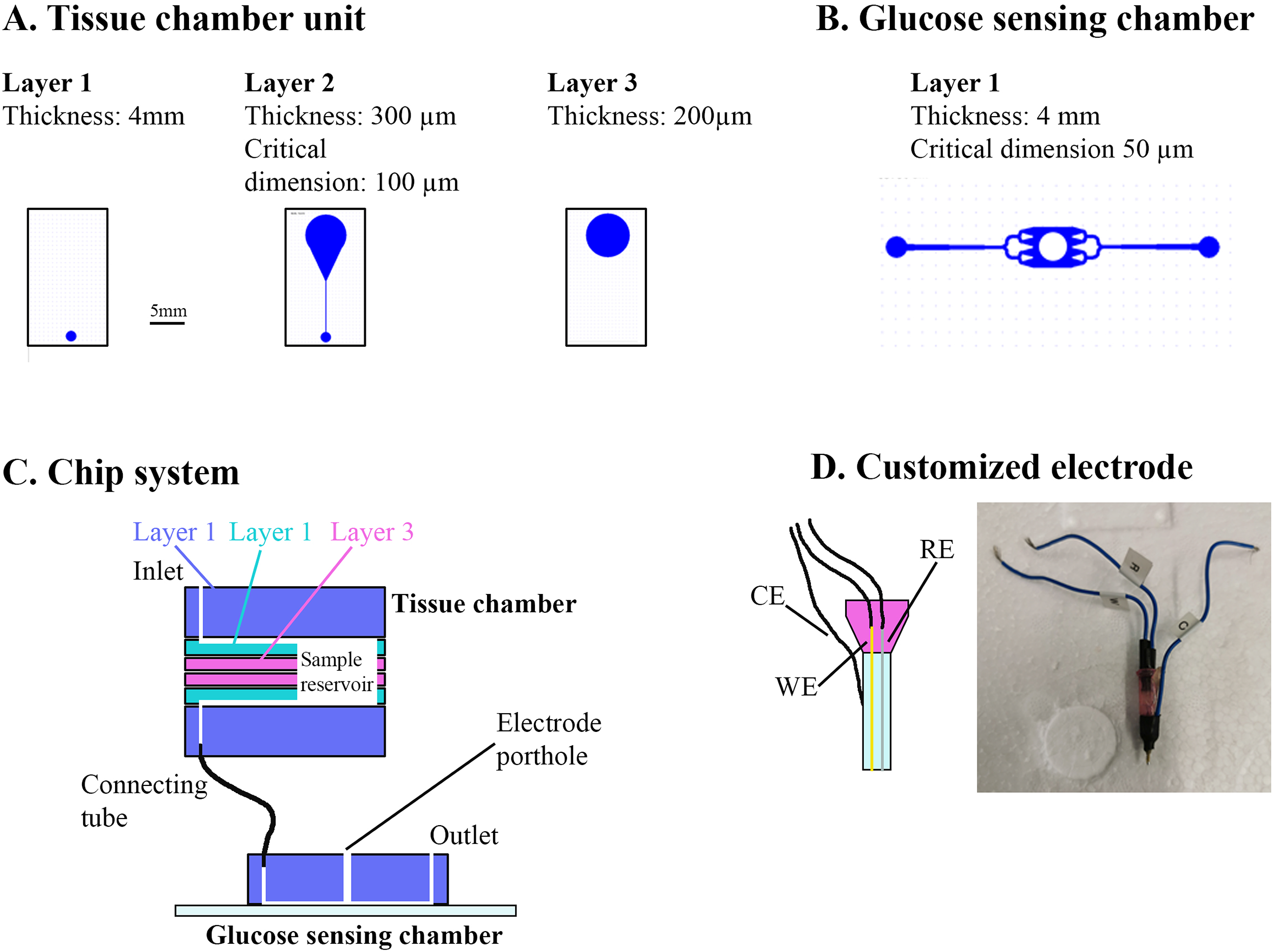
Overview of muscle chip for glucose sensing. A) Overview of the different PDMS layers for the tissue chamber unit. Scale bar = 5 mm. B) Microfluidic system for the glucose sensing chamber. The electrode was placed in the center of the system in the punched whole. C) Overview of the muscle chip system showing the various layers of the tissue chamber as well as the connection to the glucose sensing chamber. D) Schematic drawing and picture of the customized electrode fabricated to sense glucose based on glucose oxidase. RE = reference electrode, CE = counter electrode, WE = working electrode.

In parallel, FDB fibers were isolated as described but cultured on 4 mm paper patches (filter 114, Whatman) by diluting the fiber solution 5:1 in ECM gel and adding 50 µl to each patch. Two hours prior to experiment the fibers were starved from serum and glucose. Upon experiment the microfluidic system was assembled. First the electrode was inserted into the detection chamber and calibrated by perfusing solutions with known glucose concentrations through the whole system. Next the fiber-containing paper patch were inserted into the tissue chamber and perifused with serum free DMEM containing 4 mM glucose for 3-5 minutes. Next, the glucose concentration in the perfusate was monitored during basal and insulin-stimulated conditions and Δglucose was calculated as previously described (81). Colchicine and Noco treatment for 2h was achieved by preincubating the paper patches containing fibers in colchicine and Noco and keeping the drugs in the perfusate at all times after the assembly of the tissue chamber. Noco treatment for 5 min was achieved by switching the perfusate to one containing Noco 5 minutes prior to insulin stimulation.

### Western blotting

Samples were lysed in lysis buffer (50 mM Tris base, 150 mM NaCl, 1mM EDTA, 1 mM EGTA, 50 mM sodium fluoride, 5 mM pyrophosphate, 2 mM sodium orthovanadate, 1 mM dithiothreitol, 1 mM benzamidine, 0.5% protease inhibitor cocktail (Sigma P8340), 20% NP-40, pH 7.4) before processing as previously described (82). The following antibodies were used: phospho (p)-Akt Thr308 (9275, CST), Akt (9272, CST), p-TBC1D4 Thr642 (4288, CST) or TBC1D4 (ab189890, Abcam). Coomassie stain was used as a loading control.

### Immunolabelling

Human fiber bundles were teased into individual fibers and transferred to wells in a 24-well plate containing PBS using fine forceps. FDB fibers were similarly incubated in PBS. Fibers were washed 3×10 min in PBS and incubated in blocking buffer (1% bovine serum albumin (Merck), 5% goat serum (16210-064, Gibco), 0.1% Na Azide (247-852, Merck), 0.04% Saponin (27534-187, VWR) for 1 hour. The muscle fibers were then incubated in blocking buffer containing primary antibodies overnight at 4°. The next day the fibers were washed 3×10 min in PBS containing 0.04% saponin and incubated in blocking buffer with Alexa 488 anti rabbit or Alexa 568 anti rabbit or anti mouse (Invitrogen) for 2 hours. Finally, the fibers were washed 3×10 min in PBS and mounted on glass slides in Vectashield (H-1000, Vector Laboratories) or imaged directly from the glass bottom dish. The following antibodies raised in rabbit: GLUT4 (PA5-23052, Invitrogen), Detyrosinated α-tubulin (AB48389, Abcam), rabbit Syntaxin6 (110 062, Synaptic Systems), or in mouse: GLUT4 (MAB8654, R&D Systems), α-tubulin (T9026, Merck).

### Image acquisition and processing

Imaging was performed using the following systems: Zeiss 710, 780, 900, 980 or Elyra PS.1. Confocal imaging was performed using a Zeiss Plan-Apochromat 63x 1.4 NA objective. Laser source was an Argon laser emitting excitation light at 488 nm (25mW) and helium neon laser emitting excitation light at 543 nm (1.2mW), assembled by Zeiss. Emission light was collected by PMTs with matching beam splitters by Zeiss. The different channels were acquired sequentially. All live-imaging was performed in an integrated incubator at 37° in 5% CO_2_ and the fibers were kept in α-MEM containing drug/hormone as described. Specific imaging details for time series are provided in figure legends. Time series with 1 image per second for 60 seconds were obtained for mCherry-Tubulin and GLUT4-GFP dual color imaging. EB3-GFP time series were 1 image per second for 30 or 60 seconds. GLUT4-GFP imaging were time series of 30 to 300 seconds with 0.1 to 1 image per second. At all times pixel size was kept at ≤90×90 nm. The pixel dwell time was 1.27 µs. To make dynamics perceptible, color coded projections were generated as described. From this, moving objects appear rainbow-colored whereas static objects appear white.

Structured Illumination Imaging (SIM) was performed using an Elyra PS.1 system (Zeiss), with a Zeiss Plan-Apochromat 63x 1.4 NA objective and a 1.6x tube lens. The system was driven by Zen Black 2.3 SP1 from Zeiss which automatically assigns a diffraction pattern for each used wavelength (namely 28 um for 488 nm, and 34 um for 561 nm). Laser source was diode lasers, emitting excitation light at 488 nm (200 mW) and 561 nm (200 mW), assembled by Zeiss. Emission light was collected by a PCO.edge 5.5 sCMOS camera (PCO, Kelheim, Germany) with matching beam splitters by Zeiss. The different channels were acquired sequentially. Settings for image collection aimed at obtaining a dynamic range >5000 grayscale levels, and bleaching was assessed to be <20% of the dynamic range across the imaging sequences. 3D stacks were acquired at 100 nm steps by using a PI E-655 Z-piezo drive by Physik Instrumente (Karlsruhe, Germany).

Unless otherwise noted images shown are single frames. For visualization purposes only, some images were cropped and contrast or levels were linearly adjusted. Images were processed using ImageJ (83) and Adobe Photoshop 21.2.1 (Adobe).

### 3D-SIM image reconstruction

The Structured Illumination image processing module built in Zen Black 2.3, keeping a theoretical PSF and the same noise filter parameter (−5.5) for all the processed images was used for reconstruction. The resulting super-resolution images were kept raw scaling and were baseline shifted. Super-resolution image quality was assessed by applying FFT to the reconstructed images, compared to FFT of the widefield acquisition. System performance was assessed by using an Argolight SIM patterned standard sample (Argolight, Pessac, France), obtaining resolutions ∼120 nm consistently, and PSF was evaluated using 100 nm tetrapeck beads from Zeiss.

### Live-imaging movies

Representative movies were generated from the live-imaging time series in 10 frames per second. For GLUT4-GFP and mCherry-Tubulin dual color time series (movie 1), the movie was generated by merging the two channels with the GLUT4-GFP channel being green and the tubulin channel being magenta. To facilitate visualization, the single-color time series (movie 2-4) were generated by removing every other frame and switching the colors from green to red between remaining frames.

### Depletion and relocalization analysis

Using the particle analysis tool in ImageJ (83), GLUT4-GFP vesicles (sized >0.02 µm^2^, circularity between 0 and 1) from background and threshold adjusted 8-bit images were identified and vesicle areas were determined. From this small vesicles <0.04 µm^2^ were counted and related to the total number of vesicles as a reference for overexpression. These vesicles were sized the smallest resolvable and up to ∼225 nm in diameter. We analyzed this fraction since insulin induces membrane insertion of the small 50-70 nm GSVs (84) which are expected to be part of this fraction. For relocalization analysis vesicles were identified similar as for the depletion analysis. The individual vesicles were allocated into one of the following three groups: Large vesicles (>4 µm^2^), medium sized vesicles (between 0.4 and 4 µm^2^) and small vesicles (<0.4 µm^2^).

### Polymerization rate analysis

Via calponin homology domains EB3 proteins interact with tubulin at the microtubule tip and can thus be used to identify polymerizing microtubules (35). On 8-bit threshold-adjusted images, the tip of polymerizing microtubules were identified as a 0.08-0.2 µm^2^ region with a circularity between 0.2 and 1 with accumulated EB3-GFP signal. Based on these criteria the average number of polymerizing microtubules per image in a 30 or 60 second time series with an image every second were calculated.

### Tracking analysis

was performed on 8-bit images using the ImageJ plug in TrackMate. Threshold was adjusted in TrackMate and tracking settings was adjusted to maximize fitting of the automated tracking. The following tracking settings was used, for GLUT4-GFP: LoG detector, 0.5 µm blob diameter estimate, LAP tracker, 1.5 µm gap-closing and max distance and a maximum of 1 frame gap. Settings were similar for EB3-GFP except 2 µm was used as gap-closing distance, max distance was 1 µm and max frame gap was 2. Tracks with a <1.5 µm displacement was considered not to be microtubule-based GLUT4 movement or microtubule polymerization and not included in further analysis.

### Statistical analyses

Results are shown as mean, mean with individual values or mean ± SD. Statistical testing was performed using t-test or one- or two-way ANOVA as described in the figure legends. Tukey’s post hoc test was performed following ANOVA. The significance level was set at p<0.05.

## Data availability

All data generated or analyzed during this study are included in the manuscript and supporting files.

## Acknowledgements

We acknowledge the Core Facility for Integrated Microscopy, Faculty of Health and Medical Sciences, University of Copenhagen and especially Pablo Hernández-Veras for his guidance with the SIM imaging. We acknowledge Prof. Johan Auwerx (École Polytechnique Fédérale de Lausanne) for help with the mouse work. Finally we also acknowledge, from the August Krogh Section for Molecular Physiology Section at University of Copenhagen Bente Kiens and Erik A. Richter for the expertise and help with conducting the human study.

## Author contributions

JRK and TEJ conceived the study. JRK, CHO, ZL, DES, JRH, JFPW performed the human experiment. JRK performed the rodent experiments with help from KWP, CHO, ZL, SHR, MW and TEJ. JRK, KWP and NDL performed the *in vitro* cell experiments. JRK developed the glucose sensing muscle chip with guidance from RT and MAG. JRK performed the imaging and image analyses. JRK and TEJ wrote the manuscript and all co-authors commented on the draft and approved the final version. JRK and TEJ are the guarantors of this work. As such, they have full access to all data in the study and take full responsibility for the integrity of the data and the accuracy of the data analyses.

---

**Figure 1 – figure supplement 1:**
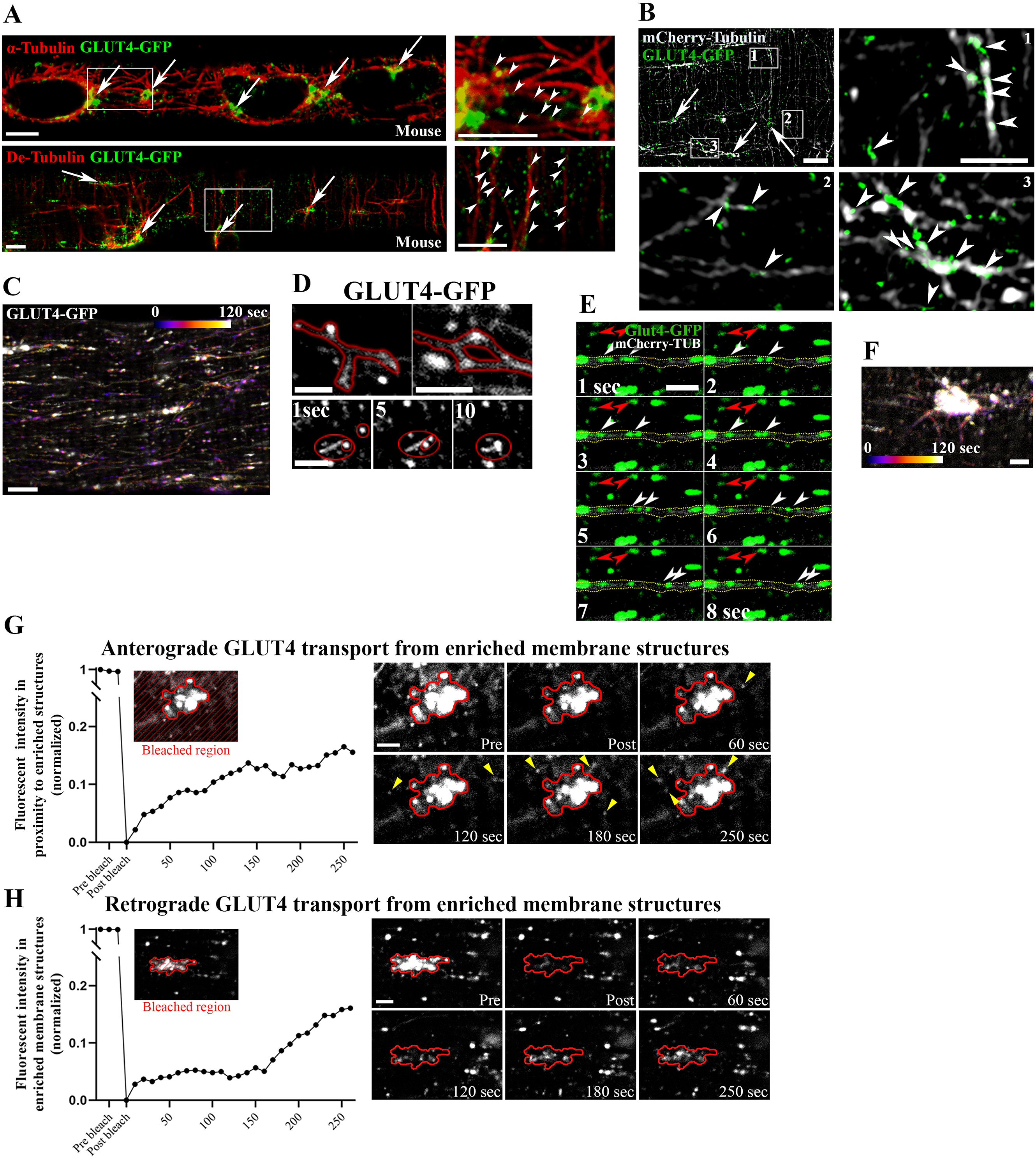
GLUT4 undergoes budding and fusion on the microtubules and moves bi-directionally to and from the microtubule nucleation sites. A) Confocal imaging of α-tubulin (top panel), detyrosinated tubulin (De-Tubulin, bottom panel) and GLUT4-GFP in flexor digitorum brevis (FDB) fibers from muscles expressing GLUT4-GFP. B) Structured illumination microscopy (SIM) imaging of FDB fiber expressing GLUT4-GFP and mCherry-Tubulin. In A and B, arrows mark GLUT4-GFP localized at microtubule nucleation sites and arrowheads indicate GLUT4-GFP on the microtubule filaments. C) Color-coded projection (first image blue, last image white as indicated) from 120 second live-imaging of FDB fiber expressing GLUT4-GFP. D) Live-imaging indicating spherical and tubular GLUT4-GFP vesicles/structures undergoing fusion and fission. E) Time-montage of GLUT4-GFP and mCherry-Tubulin from the recording shown in figure 1D. White arrowheads mark GLUT4-GFP moving on the microtubule filament incorporated with mCherry-Tubulin. Red arrowheads mark static GLUT4-GFP localized away from the microtubule filament. F) Color-coded projection (first image blue, last image white as indicated) from 120 seconds live-imaging of GLUT4-GFP-enriched structures at the microtubule nucleation sites in FDB fiber. Directionality of the GLUT4 trafficking is visible in movie 2. G) Quantified graph (left) and time series images (right) of GLUT4-GFP fluorescence in region surrounding microtubule nucleation sites (outside red outline), before and after photobleaching of the surrounding region to estimate plus-end-directed GLUT4 transport. Yellow arrowheads indicate presence of new GLUT4 structures. H) Quantified graph (left) and time series images (right) of GLUT4-GFP fluorescence at microtubule nucleation site (encircled in red), before and after photobleaching of the central nucleation site region to estimate minus-end-directed GLUT4 transport. Time series images shown are representative of fibers from 3 different mice. Images are representative of at least 5 fibers from ≥3 different mice. Scale bar = 5 µm (A-C) and 2 µm (D-H and inserts in B). **Figure 1 – figure supplement 2: GLUT4 travelled on microtubules** Movie of live FDB fibers electroporated with GLUT4-GFP (green) and mCherry-Tubulin (magenta) 6 days earlier. The movie is played in 10 frames per second with one frame representing 1 second. White arrows mark areas for frequent GLUT4-travelling. Scale bar = 5 μm. **Figure 1 – figure supplement 3: GLUT4 travelled to and from GLUT4-enriched regions at microtubule nucleation sites.** Movie of GLUT4-GFP at GLUT4 enriched/large structure in live FDB fibers showing spherical and tubular GLUT4-GFP structures that fuse and bud off from the GLUT4-enriched area. To facilitate visualization, the movie was generated so moving GLUT4 appears green-red flashing, whereas static GLUT4 appears yellow as described in the methods section. The movie is played in 10 frames per second with one frame representing 4 seconds. Scale bar = 5 μm.

**Figure 2 – figure supplement 1:**
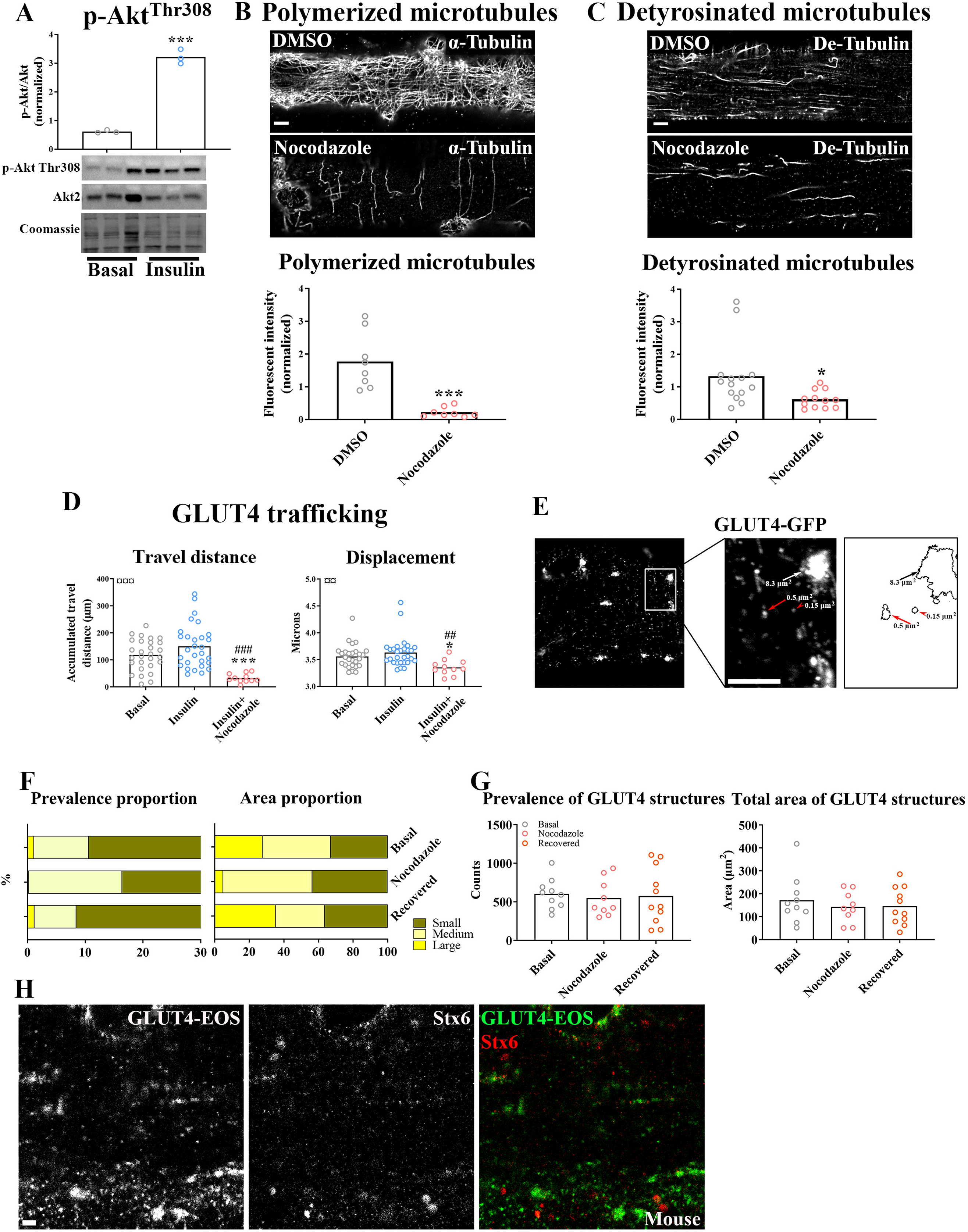
GLUT4 trafficking and localization was dependent on an intact microtubule network. Phosphorylation of Akt Thr308 in GLUT4-GFP expressing flexor digitorum brevis (FDB) fibers treated ± 30 nM insulin (INS) for 15 minutes. B) Polymerized microtubules in GLUT4-GFP expressing fibers treated ± Nocodazole (Noco, 13 µM) for 4 hours and stained with α-Tubulin. C) Fibers as in B stained for detyrosinated tubulin (De-Tubulin). D) Microtubule-based GLUT4-GFP trafficking quantified as the travelled distance and the lateral displacement of particles within the dynamic fraction in muscle fibers treated ± Noco (13 µM) for 4 hours and ± INS (30 nM) for 15-30 minutes. E) Identification of GLUT4-containing structures in GLUT4-GFP expressing FDB fibers. Black/white arrows mark large (>4 µm^2^) structures indicative of GLUT4 clusters at the microtubule nucleation sites. Red arrow marks a structure sized between 0.4 and 4 µm^2^ and categorized as a medium size structure. Red arrowhead marks a structure of <0.4 µm^2^) which is categorized as a small structure. F) Bar graphs showing the relative prevalence and area in the different membrane size categories in fibers ± microtubule network disruption by addition of Noco (13 µM) 15 hours prior to imaging or 9 hours after Noco removal. G) Total number and area of vesicles in fibers as in F. H) GLUT4-EOS and Syntaxin6 (Stx6) in mouse FDB fibers. For A, n=3 mice, for B-C n ≥ 8 muscle fibers from 3 different mice. For D, n ≥ 14 muscle fibers from 5 different mice. For F-G, n = 9-11 muscle fibers from 3 different mice. For H, n = 6-7 fibers from 2 different mice. Data are presented as mean with individual data points. */*** p<0.05/0.001 different from Basal/DMSO. ##/### p<0.01/0.001 different from INS. ¤¤/¤¤¤ p<0.01/0.001 ANOVA effect **Figure 2 – figure supplement 2: GLUT4 movement was disrupted by nocodazole treatment** Movie of live FDB fibers electroporated with GLUT4-GFP 6 days earlier. To facilitate visualization, the movie was generated so moving GLUT4 appears green-red flashing while static GLUT4 appears yellow, as described in methods section. The left panel shows fiber in the basal state, the middle panel shows fiber treated with insulin for 15 min and the right panel shows fiber treated with insulin for 15 min after a prior treatment with 4 µg/ml nocodazole for four hours. The movie is played in 10 frames per second with one frame representing 4 seconds. Scale bar = 5 μm. **Figure 2 – source data 1: Data used for quantification of GLUT4 trafficking and localization** **Figure 2 – figure supplement 1 - source data 1: Data used for quantification of GLUT4 trafficking and localization**

**Figure 3 – figure supplement 1:**
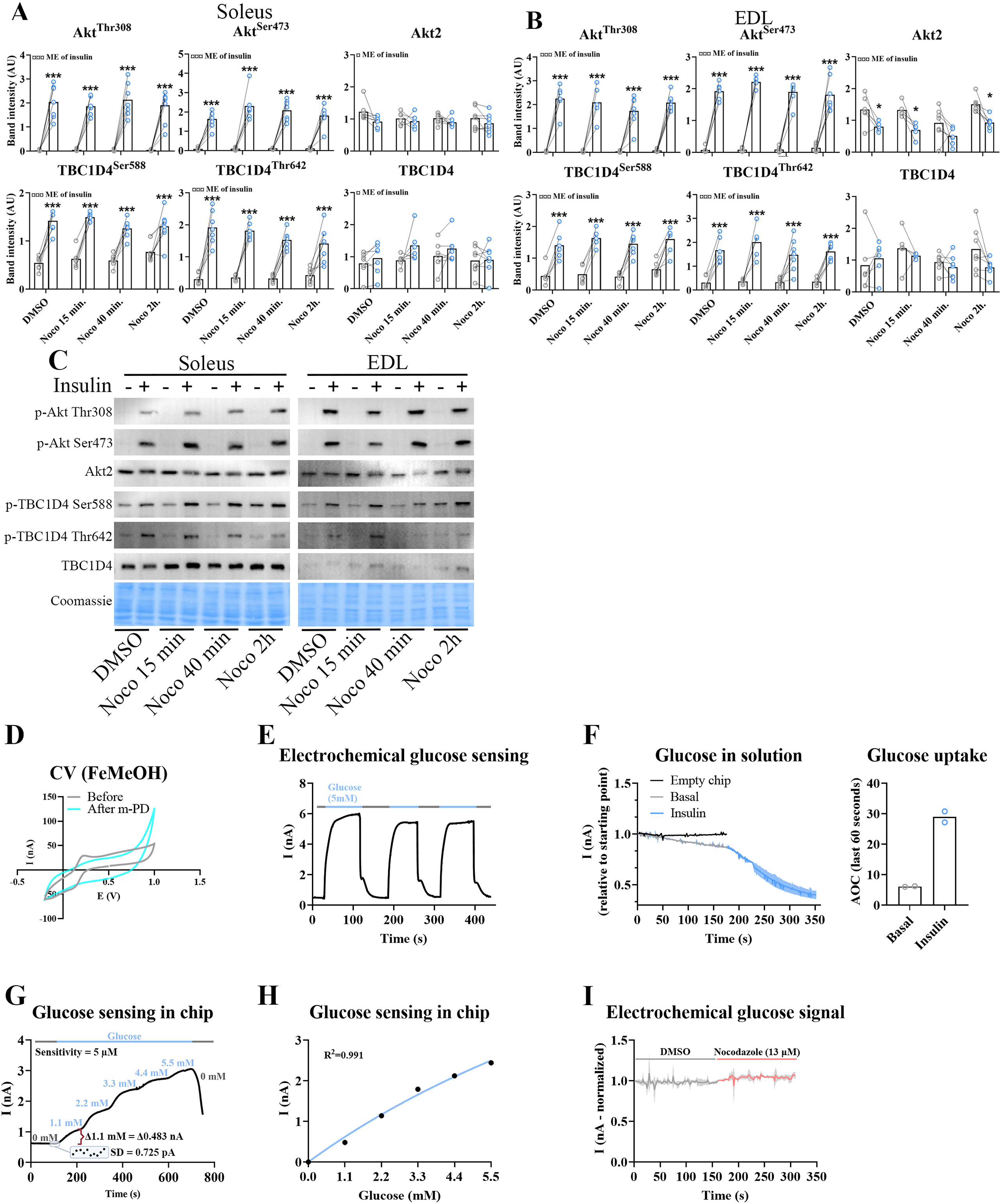
Time-dependent effect of microtubule disruption on insulin-induced muscle glucose uptake. A-B) Quantification of protein expression in soleus (A) and EDL (B) muscles in the basal and insulin-stimulated state ± Nocodazole (Noco, 13 µM) as indicated. C) Representative immunoblots of the proteins quantified in A-B. D) Deposition of poly-(m-phenylenediamine) (m-PD) film as exclusion membrane to improve sensor selectivity demonstrated by abolished anodic signal in the voltammograms (scanning rate (SR)= 100 mV s^-1^) performed at the clean Pt surface in 1 mM FcMeOH after m-PD deposition. E) Glucose oxidase presence validation by current (nA) change upon sensor transfer to and from a glucose solution (5mM). F) Current change and calculated area over the curve (AOC) in perifusate from mouse flexor digitorum brevis (FDB) fibers before and after insulin stimulation. G) Typical calibration trace (Q = 1 μl s^-1^) when exposing glucose sensor to stepwise 1.1 mM glucose increments. Sensitivity of the sensing was calculated as 3*SD / Δ current * Δ glucose. SD was calculated from 10 baseline data points, Δ current was calculated when switching from 0 to 1.1 mM glucose as indicated by the red bracket. H) Associated calibration curve, fit with a Michaelis–Menten model. I) Current fluctuations in perifusate from FDB fibers ± Noco (13 µM). ME = main effect. For A-C n = 6-7 muscles from 6-7 mice. Data are presented as mean with individual data points. Paired observations from the same mouse are indicated by a connecting line. F and I, graphs indicate mean ± standard deviation. */*** p<0.05/0.001 different from DMSO. ¤/¤¤¤ p<0.05/0.001 ANOVA effect. **Figure 3 – source data 1: Data used for quantification of glucose uptake, polymerized microtubules and GLUT4 localization in figure 3** **Figure 3 – figure supplement 1 - source data 1: Data used for quantification of protein expression and electrochemical glucose sensing in figure 3 – figure supplement 1**

**Figure 4 figure supplement 1:**
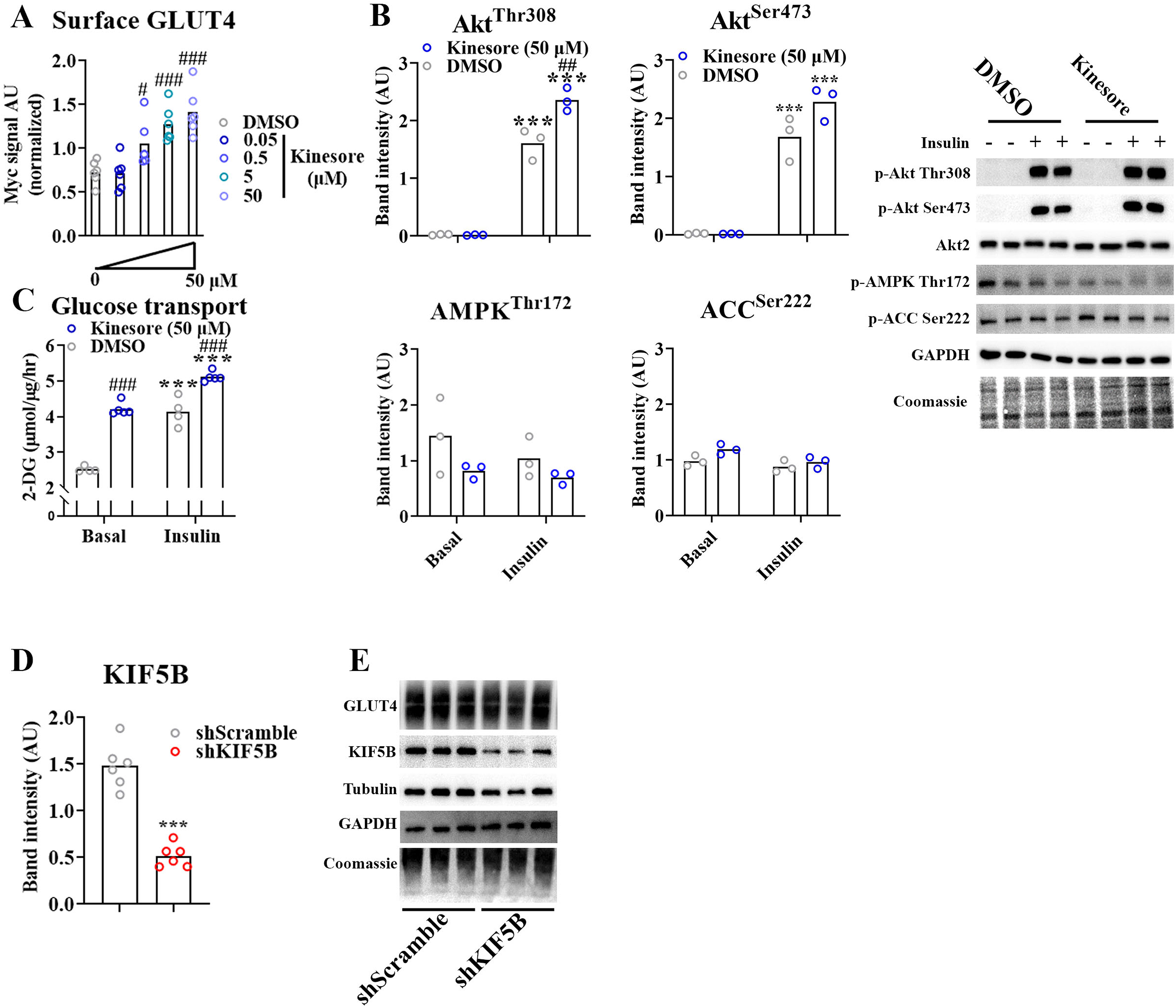
Kinesin-1 containing Kif5b regulated GLUT4 localization and translocation. *A)* Exofacial GLUT4 signal at the surface membrane of L6 myoblast serum starved for 4h and treated ± kinesore at indicated concentration for the last 2h before cell fixation. ANOVA effect (¤¤¤). B) Quantification and representative immunoblots showing protein expression and phosphorylation in basal and insulin stimulated (100 nM, 15 min) L6 myoblasts serum starved for 4h and treated ± kinesore for 2h. ANOVA main effect of insulin (Akt Thr308+Ser473 = ¤¤¤, ACC Ser222 = ¤), ANOVA main effect of Kinesore (ACC Ser222 = ¤, Akt Ser473+AMPK Thr172 p=0.07), interaction (Akt Thr308 = ¤¤, Akt Ser473 p=0.06). C) 2-deoxyglucose (2-DG) transport in L6 myoblast serum starved for 4h and treated ± kinesore (50 µM) for 2h before 20 min ± insulin (100 nM) and 5 min 2-DG accumulation. ANOVA main effect of insulin (¤¤¤) and Kinesore (¤¤¤) and interaction (¤¤). D) Quantification of KIF5B protein expression in L6 myoblasts transfected with shScramble RNA or shRNA targeting Kif5b for 72h. E) Representative immunoblots showing protein expression in L6 myoblasts as in D. A, n = 6 replicates from 2 independent experiments. B, n = 3 independent experiment. C, n = 5-6 replicates from 2 independent experiments. D-E, n = 6 replicates from 2 independent experiments. *** p<0.001 effect of insulin/shKif5b. #/##/### p<0.05/0.01/0.001 different from DMSO. ¤/¤¤/¤¤¤ p<0.05/0.01/0.001 ANOVA effect. **Figure 4 – source data 1: Data used for quantification of GLUT4 localization and GLUT4 surface content in figure 4** **Figure 4 – figure supplement 1 - source data 1: Data used for quantification of GLUT4 surface content, protein expression and glucose uptake in figure 4 – figure supplement 1**

**Figure 5 – figure supplement 1:**
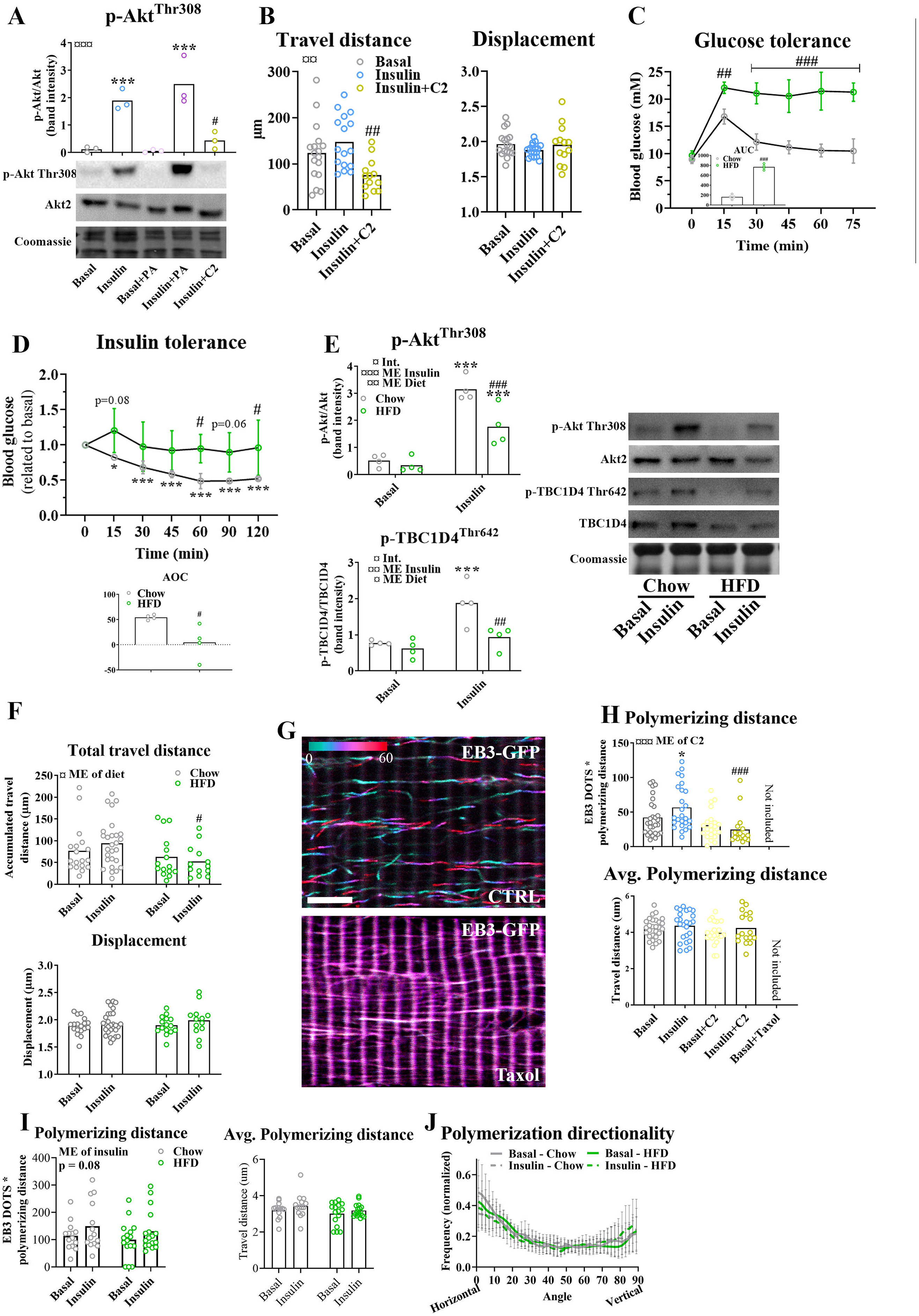
Insulin resistance impairs microtubule-based GLUT4 trafficking. A) Phosphorylation of Akt^Thr308^ in GLUT4-GFP expressing flexor digitorum brevis (FDB) muscle fibers cultured ± 0.5 mM palmitic acid (PA) for 24 hours or 50 µM C2 ceramide (C2) for 2 hours prior to 15 minutes ± 30 nM insulin (INS) stimulation. B) Quantified microtubule-based GLUT4 trafficking assessed as the travelled distance and the displacement within the dynamic fraction of the GLUT4-GFP in basal, insulin (INS, 30 nM) or insulin and C2 Ceramide (INS+C2, 30 nM+50 µM)-treated FDB muscle fibers. C-D) Glucose and insulin tolerance tests of mice fed a control chow or a high fat diet (HFD) after intraperitoneal injection of glucose (2 g*kg^-1^ body weight) or insulin (0.5 U*kg^-1^ body weight). AUC/AOC = area under/over the curve. ANOVA main effect of diet (¤¤¤/¤, GTT/ITT), and time (¤¤¤) and interaction (¤¤¤/¤¤, GTT/ITT). E) Phosphorylation of Akt^Thr308^ and TBC1D4^Thr642^ in isolated FDB fibers from muscles from chow or HFD fed mice treated ± INS (30 nM) for 15 min. F) Quantified microtubule-based GLUT4 trafficking assessed as the travelled distance and the displacement within the dynamic fraction of the GLUT4-GFP in basal or INS(30 nM)-stimulated FDB muscle fibers isolated from chow or HFD-fed mice. G) Color coded projection ((first image cyan, last image red as indicated by color code bar) of 60 sec. live-imaging of FDB fibers expressing EB3-GFP and treated without (CTRL) or with 10 µM paclitaxel (Taxol) for 2 hours. Scale bar = 5 µm. EB3-GFP dynamics are also illustrated in movie 4. H) Quantification of the total and average polymerizing distance of EB3-GFP containing microtubules in fibers incubated ± 50 µM C2 ceramide (C2) or 10 µM Taxol for 2 hours prior to 15-30 minutes of INS (30 nM) stimulation. I) Quantification of the total and average polymerizing distance of EB3-GFP containing microtubules in fibers isolated from chow or HFD fed mice and stimulated ± INS (30 nM) for 15 minutes. J) Quantification of microtubule polymerization directionality based on EB3-GFP dynamics recording as in I. For A, C-E n=3-4 mice, for B, F and H-J n ≥ 13 muscle fibers from 3-4 mice. G) representative of >10 fibers from 2 different mice. Taxol treated muscle fibers were only used as a reference and not included in the statistical analysis. NA = not analysed. Data are presented as mean with individual data points or mean±SD. **/*** p<0.01/0.001 different from basal, #/##/### p<0.05/0.01/0.001 different from insulin (A-B) or different from corresponding group in chow fed mice (C-F, H). ¤/¤¤/¤¤¤ p<0.05/0.01/0.001 ANOVA main effect (ME). **Figure 5 – figure supplement 2:** ***Polymerizing microtubules in muscle fibers*** Movie of polymerizing microtubule (MT) tips detected by EB3-GFP in live FDB fibers electroporated with EB3-GFP 6 days earlier. To facilitate visualization, the movie was generated so moving MT tips (EB3 dots) appear green-red flashing whereas static EB3 appear yellow, as described in methods section. The movie is played in 10 frames per second with one frame representing 4 seconds. Scale bar = 5 μm. **Figure 5 – source data 1: Data used for quantification of GLUT4 trafficking and microtubule polymerization in figure 5** **Figure 5 – figure supplement 1 - source data 1: Data used for quantification of protein expression, GLUT4 trafficking, glucose and insulin tolerance, microtubule polymerization and polymerization directionality in figure 5 – figure supplement 1**

